# Olfactory learning in *Drosophila* requires O-GlcNAcylation of mushroom body ribosomal subunits

**DOI:** 10.1101/2023.09.14.557796

**Authors:** Haibin Yu, Dandan Liu, Yaowen Zhang, Ruijun Tang, Xunan Fan, Song Mao, Lu Lv, Fang Chen, Hongtao Qin, Zhuohua Zhang, Daan M.F. van Aalten, Bing Yang, Kai Yuan

## Abstract

O-GlcNAcylation is a dynamic post-translational modification that diversifies the proteome. Its dysregulation is associated with neurological disorders that impair cognitive function, and yet identification of phenotype-relevant candidate substrates in a brain-region specific manner remains unfeasible. By combining an O-GlcNAc binding activity derived from *Clostridium perfringens* OGA (*Cp*OGA) with TurboID proximity labeling in *Drosophila*, we developed an O-GlcNAcylation profiling tool that translates O-GlcNAc modification into biotin conjugation for tissue-specific candidate substrates enrichment. We mapped the O-GlcNAc interactome in major brain regions of *Drosophila* and found that components of the translational machinery, including many ribosomal subunits, were abundantly O-GlcNAcylated in the mushroom body, the computational center of the *Drosophila* brain. Hypo-O-GlcNAcylation induced by ectopic expression of active *Cp*OGA in the mushroom body decreased local ribosomal activity, leading to olfactory learning deficits that could be rescued by increasing ribosome biogenesis. Our study reveals that O-GlcNAcylation contributes to the links between protein synthesis and cognitive function in the brain learning center, and provides a useful tool for future dissection of tissue-specific functions of O-GlcNAcylation in *Drosophila*.

## Introduction

Protein O-GlcNAcylation is a ubiquitous post-translational modification that occurs on thousands of nuclear and cytoplasmic proteins, conveying various stimuli or stressors such as fluctuating nutrient levels to distinct cellular processes^1-3^. It involves reversible attachment of β-*N*-acetylglucosamine (GlcNAc) to the hydroxyl group of serine and threonine residues of protein substrates, catalyzed by a pair of evolutionarily conserved enzymes, O-GlcNAc transferase (OGT) and O-GlcNAcase (OGA)^4^. As a monosaccharide modification, the addition and removal of O-GlcNAc moiety are dynamic, with cycling rates ranging from several minutes to the lifetime of a protein^5,6^. By modifying different protein substrates, O-GlcNAcylation exerts critical regulatory functions in a wide range of basic cellular processes, including transcription, translation, and protein homeostasis^1,7^. O-GlcNAcylation is ubiquitously distributed but more abundant in some tissues, such as the brain ^8,9^. Given its enrichment in brain tissues and essential biological functions, it is not surprising that O-GlcNAc cycling is required for the development and functions of central nervous system^2,10,11^, and its dysregulation is linked to numerous neurological disorders^7,10,12,13^.

O-GlcNAc homeostasis appears to be required for proper cognitive function, although the molecular connections between the dysregulated O-GlcNAcome and cognitive impairment are not fully understood. Hypomorphic mutations of *OGT* are implicated in an X-linked intellectual disability syndrome^14-18^, a severe neurodevelopmental disorder now termed *OGT*-associated Congenital Disorder of Glycosylation (OGT-CDG)^19^. *Drosophila* models of OGT-CDG that carry the equivalent human disease-related *OGT* missense mutations manifest deficits in sleep and habituation, an evolutionarily conserved form of non-associative learning ^20^. Our recent work has shown that decreased O-GlcNAcylation level in *Drosophila*, induced through overexpression of a bacterial OGA from *Clostridium perfringens* (*Cp*OGA), leads to a deficit of associative olfactory learning. More interestingly, ectopic expression of *Cp*OGA during early embryogenesis results in reduced brain size and learning defect in adult flies, likely due to interference of the sog-Dpp signaling required for neuroectoderm specification^21^. These studies reveal that disturbed O-GlcNAc homeostasis can impact cognitive function by compromising neuronal development. On the other hand, a number of studies have revealed that impaired O-GlcNAcylation is implicated in aging-related neurodegenerative diseases such as Alzheimer’s disease (AD)^7,10,12,13,22^. In the cerebrum of AD patients, O-GlcNAcylation levels are significantly lower than that of healthy controls^23^. Increased O-GlcNAcylation levels by limiting OGA activity recovers the impaired cognitive function in AD mice models^24,25^. Interestingly, during normal aging in mice, reduction of O-GlcNAcylation levels also occurs in the hippocampus, and elevation of neuronal O-GlcNAc modification ameliorates associative learning and memory^26^. These results indicate that, in addition to its involvement in neurodevelopoment, O-GlcNAc homeostasis is also required for normal neuronal activity and cognitive function. However, the identity of key O-GlcNAc protein substrates supporting the cognitive abilities in adult brain and their spatial distribution remain largely unknown.

An obstacle for comprehensively identifying the O-GlcNAc conveyors of various cognitive functions is the lack of an effective tissue-specific O-GlcNAc profiling method. Given the structural diversity and relatively low abundance, enrichment of O-GlcNAc modified proteins is required for mass spectrometry (MS)-based profiling of O-GlcNAcylation^27^. The enrichment strategies roughly fall into two categories. One category involves direct capture of O-GlcNAcylated proteins by antibodies or lectins that recognize the GlcNAc moiety^27-33^. O-GlcNAc antibodies including RL2 and CTD110.6, as well as O-GlcNAc-binding lectins such as wheat germ agglutinin (WGA), are commonly used for enrichment. In addition, the catalytic-dead mutant of *Cp*OGA that retains the ability to recognize O-GlcNAcylated substrates was successfully repurposed to concentrate many developmental regulators from *Drosophila* embryo lysates^34^. Another category of enrichment strategies relies on chemoenzymatic or metabolic labeling^27-33^. Azido-modified intermediates, such as *N*-azidoacetylglucosamine (GlcNAz) and *N*-azidoacetylgalactosamine (GalNAz), are used to introduce specific tags (e.g. biotin) to protein substrates via Staudinger ligation or click chemistry, allowing for capture and enrichment of O-GlcNAcylated proteins. A recent study coupled the O-GlcNAc-binding lectin GafD to the proximity labeling TurboID yielding the GlycoID tool^35^, in which GafD domain recognizes O-GlcNAcylated substrates and the TurboID enzyme attaches nonhydrolyzable biotin tags to proximal proteins within approximately 10 nm radius^36^. The GlycoID tool was used to profile O-GlcNAcylation in different subcellular spaces including nucleus and cytosol^35^. These different O-GlcNAcylation profiling strategies have greatly expanded the O-GlcNAcome over the past 30 years^9,32^. However, none of them have been adopted for tissue-specific profiling of O-GlcNAcylated proteins.

Here we generated transgenic *Drosophila* lines that allow specific expression of *Cp*OGA in different brain regions. Ectopic expression of *Cp*OGA in the major learning center of *Drosophila* brain, the mushroom body, reduced local O-GlcNAcylation levels and impaired olfactory learning. We further combined a catalytically incompetent *Cp*OGA mutant (*Cp*OGA^CD^) with the proximity labeling enzyme TurboID to develop an O-GlcNAcylation profiling tool. By conditional expression of this tool to translate O-GlcNAc modification into biotin conjugation in specific brain structures, we mapped the O-GlcNAc interactomes and generated an O-GlcNAc atlas for different brain regions of *Drosophila* (tsOGA, http://kyuanlab.com/tsOGA/). Particularly, we detected abundant O-GlcNAc modifications associated with ribosomes in the mushroom body. Lowering the mushroom body O-GlcNAcylation levels reduced the ability of ribosomes to synthesize new proteins, interfering with olfactory learning, which could be reversed by increasing ribosomal biogenesis via overexpression of dMyc. We propose that compromised ribosomal activity in the brain learning center contributes to the cognitive deficits of O-GlcNAcylation insufficiency-associated neurological diseases.

## Results

### Perturbation of the mushroom body O-GlcNAcylation leads to olfactory learning deficits

We previously reported that ubiquitous expression of *Cp*OGA in *Drosophila* reduced global O-GlcNAcylation levels and resulted in impaired olfactory learning^21^. To determine which brain region was critical for this hypo-O-GlcNAcylation induced learning defect, we conditionally expressed wild type *Cp*OGA (*Cp*OGA^WT^) in different brain structures of *Drosophila*, driven by Elav-Gal4 (pan-neuron), OK107-Gal4 (mushroom body), C232-Gal4 (ellipsoid body), GMR14H04-Gal4 (antennal lobe), and GMR33H10-Gal4 (optic lobe) respectively (Figure 1A). *Cp*OGA^DM^, which carries two point-mutations (D298N and D401A) that inactivate both the catalytic and binding activities toward O-GlcNAc modification, was used as a control. We dissected brains from the adult flies and validated tissue-specific expression patterns via immunostaining. As expected, Elav-Gal4 induced *Cp*OGA^WT^ expression in the whole brain (Figure 1B), leading to decreased O-GlcNAcylation levels compared to the *Cp*OGA^DM^ (Figure 1C). Similarly, other tissue-specific Gal4 drivers activated *Cp*OGA expression in different brain structures and perturbed local O-GlcNAc modifications. For instance, OK107-Gal4 drove *Cp*OGA^WT^ expression in the mushroom body and downregulated O-GlcNAcylation levels in the Kenyon cells (Figure 1D and 1E).

**Figure 1.**
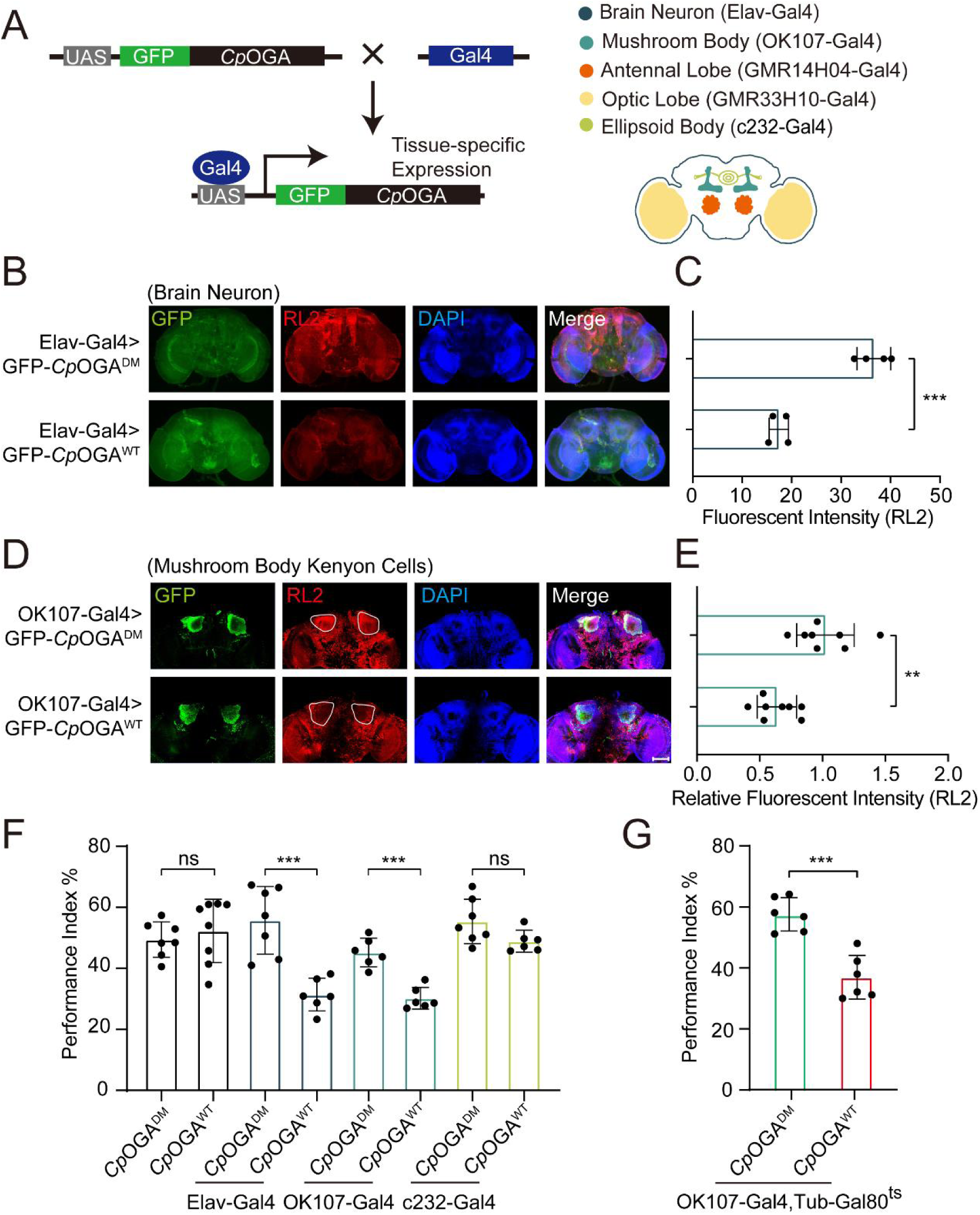
Downregulation of protein O-GlcNAcylation level in brain or mushroom body neurons affects olfactory learning of adult flies. (A) Scheme for expression of *Cp*OGA^WT^ or *Cp*OGA^DM^ in various *Drosophila* brain structures using different Gal4 drivers. (B) Immunostaining of adult *Drosophila* brains. Brains were stained with anti-O-GlcNAc antibody RL2 (red) to assess O-GlcNAcylation level, and anti-GFP (green) antibody to validate tissue-specific expression of *Cp*OGA. Nuclei were stained with DAPI (blue). Scale bar: 100 μm. (C) Quantification of fluorescent intensity of O-GlcNAc staining in *Cp*OGA^WT^ or *Cp*OGA^DM^ expressed brains. (D) Immunostaining of adult *Drosophila* brains. Outlined areas indicate the cell bodies of kenyon cells in mushroom body. Scale bar: 100 μm. (E) Quantification of relative fluorescent intensity of O-GlcNAc staining in *Cp*OGA^WT^ or *Cp*OGA^DM^ expressed brain structures. (F) A compilation of performance index in learning test of the indicated flies expressing either *Cp*OGA^WT^ or *Cp*OGA^DM^. (G) A compilation of learing performance index of flies expressing *Cp*OGA^WT^ or *Cp*OGA^DM^ only in the mushroom body at adult stage. Each datapoint represents an independent experiment with approximately 200 flies. *p* values were determined by unpaired *t*-test, and the stars indicate significant differences (\*\*\**p* < 0.001, \*\**p* < 0.01 and ns, not significant, *p* ≥ 0.05). Error bars represent SD. **Figure 1—source data 1.** Excel spreadsheet containing source data used to generate Figures 1C-G.

We then evaluated the cognitive ability of these flies using olfactory learning assay as previously reported^37-39^. To rule out the possibility that overexpression of *Cp*OGA^WT^ or *Cp*OGA^DM^ differentially disrupted odor preference, we tested their olfactory acuity toward either 4-methylcyclohexanol (MCH) or octanol (OCT) using air as a control. Tissue-specific expression of *Cp*OGA^WT^ or *Cp*OGA^DM^ in the antennal and optic lobes generated differences in susceptibility, and these flies were not included in subsequent olfactory learning tests (Figure S1A and S1B). Flies expressing *Cp*OGA^WT^ or *Cp*OGA^DM^ in brain neurons, mushroom body, or ellipsoid body were trained to associate electric shock punishment with an air current containing MCH or OCT, and then tested for the ability to remember the electric shock-associated odor using a T-maze apparatus (Figure S1C). Compared to *Cp*OGA^DM^, conditional expression of *Cp*OGA^WT^ in brain neurons or mushroom body compromised the ability to establish the association between odor and electric shock (Figure 1F), suggesting that decreased O-GlcNAcylation levels in these brain regions resulted in a deficit in olfactory learning. In contrast, flies expressing *Cp*OGA^WT^ or *Cp*OGA^DM^ in the ellipsoid body, as well as the control flies without a Gal4 driver, showed no statistical difference in the learning performance (Figure 1F). Ectopic expression of *Cp*OGA^WT^ in the mushroom body driven by OK107-Gal4 impacted neuronal development during the larval stages. To directly investigate whether perturbation of O-GlcNAcylation compromised neuronal function in adult flies, we used the temperature-sensitive Gal80 (Gal80^ts^) to restrict *Cp*OGA expression until adulthood (Figure S1D). This temporally controlled expression of *Cp*OGA^WT^ specifically in the adult mushroom body did not affect the odor acuity but significantly disrupted olfactory learning relative to *Cp*OGA^DM^ control (Figure 1G, S1A, and S1B). These results suggested that proper O-GlcNAcylation homeostasis is essential for the mushroom body function.

### O-GlcNAcylation profiling through CpOGA proximity labeling

The mushroom body is known to be the associative learning center in *Drosophila* brain^40,41^. Having discovered that O-GlcNAcylation homeostasis in the mushroom body was critical for olfactory learning, we developed an O-GlcNAc profiling method that allows identification of candidate O-GlcNAcylated protein substrates in this brain region. Mutation of the catalytic residue Asp298 to Asn (D298N) of *Cp*OGA (*Cp*OGA^CD^) inactivates the enzymatic activity but retains its ability to bind O-GlcNAcylated peptides. Taking advantage of this property, far western, gel electrophoresis, proximity ligation, and imaging methods have been developed^34,42-45^, and immobilized *Cp*OGA^CD^ has been successfully used to enrich O-GlcNAcylated substrates *in vitro*^34^. We linked this O-GlcNAc binding activity of *Cp*OGA^CD^ with TurboID, a biotin ligase that catalyzes biotinylation of adjacent proteins^36^, to tag the O-GlcNAcylated proteins with biotin for subsequent enrichment and Mass Spectrometry (MS) identification (Figure 2A and 2B). *Cp*OGA^DM^ was adopted as a control to eliminate O-GlcNAc independent protein-protein interactions (Figure 2B). Once induced by different tissue-specific drivers, this tool could tag and enrich O-GlcNAc substrates and their interactors in a tissue-specific manner, as endogenous protein biotinylation level is low in most organisms including *Drosophila*.

**Figure 2.**
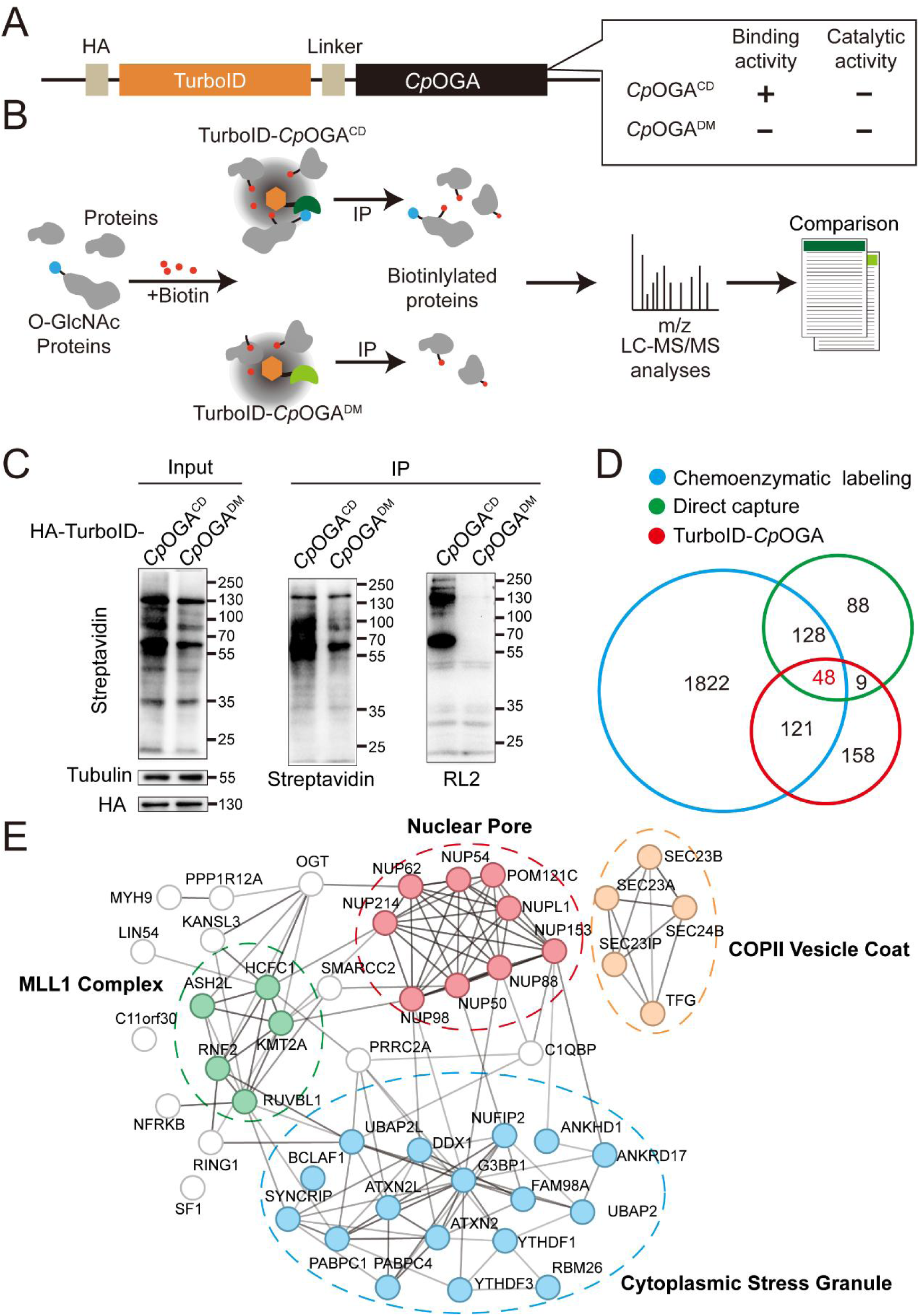
TurboID-*Cp*OGA^CD^ mediated proximity labeling of O-GlcNAc substrates in HEK293T cells. (A) Diagram of the constructs used for the expression of TurboID-*Cp*OGA^CD/DM^. (B) Schematic representation of TurboID-*Cp*OGA^CD^ based profiling strategy. In the presence of biotin, TurboID biotinylates the *Cp*OGA^CD^-bound O-GlcNAc proteins, which can be further purified by streptavidin pull-down for mass spectrometry (MS) identification. TurboID-*Cp*OGA^DM^ is used as negative control for O-GlcNAc independent protein-protein interactions. (C) Immunoprecipitation of biotinylated proteins from HEK293T cell lysates using streptavidin-magnetic beads. Biotinylation was detected by immunoblotting with streptavidin-HRP, and O-GlcNAcylation with anti-O-GlcNAc antibody (RL2). The expression of TurboID-*Cp*OGA^CD/DM^ was verified by anti-HA immunoblotting. (D) Venn diagram showing the overlap of potentially O-GlcNAcylated proteins identified with TurboID-*Cp*OGA versus that with another two commonly used methods. (E) STRING visualization of protein-protein interaction network of the 48 highly-confident O-GlcNAc substrates in HEK293T cells. **Figure 2—source data 1.** Raw data of all western blots from Figure 2. **Figure 2—source data 2.** Complete and uncropped membranes of all western blots from Figure 2. **Figure 2—source data 3.** Excel spreadsheet containing source data used to generate Figures 2C.

As proof of concept, we generated stable HEK293T cells expressing TurboID-*Cp*OGA^CD^ or its reference construct TurboID-*Cp*OGA^DM^. To characterize labeling activity, treatment with 10 μM or 100 μM biotin from an aqueous stock was first applied on these cells for 60 min, and the cell lysates were subject to western blot with streptavidin-HRP (Figure S2A). 10 μM biotin treatment yielded robust biotinylation of proteins, and this concentration was selected for subsequent experiments on cultured cells. To determine optimal incubation time, the cells were treated with 10 μM biotin from 15 to 180 min. Significant time-dependent labeling activity of proteins was observed, and 120 min was selected because it generated strong biotinylation in cells expressing *Cp*OGA^CD^ compared to the *Cp*OGA^DM^ control (Figure S2B). We validated whether fluctuation in O-GlcNAcylation could be translated into biotinylation alterations. To this end, the cells were first treated with OGA inhibitor Thiamet-G or OGT inhibitor OSMI for 6 h followed by biotin incubation. Thiamet-G increased global O-GlcNAcylation levels, and the overall biotinylation was consistently upregulated. Conversely, OSMI treatment decreased both O-GlcNAcylation and biotinylation in the cell lysates, suggesting that TurboID-*Cp*OGA^CD^ effectively translates O-GlcNAc modification into biotin conjugation (Figure S2C and S2D).

To test whether TurboID-*Cp*OGA^CD^ could be used to enrich and identify O-GlcNAcylated substrates, we performed immunoprecipitation with streptavidin magnetic beads from equal amount of cell lysates expressing either TurboID-*Cp*OGA^CD^ or TurboID-*Cp*OGA^DM^ after biotin incubation (Figure 2C). TurboID-*Cp*OGA^CD^ labeled more proteins with biotin in the input compared to TurboID-*Cp*OGA^DM^, and consistently, more biotinylated proteins were immunoprecipitated. Importantly, western blot with anti-O-GlcNAc antibody RL2 detected strong O-GlcNAcylation signals in immunoprecipitants from the cells expressing TurboID-*Cp*OGA^CD^ but not TurboID-*Cp*OGA^DM^, indicating successful enrichment of O-GlcNAc substrates using the biotin tags (Figure 2C). We scaled up the experiments and carried out MS analysis on the immunoprecipitants. Proteins that were selectively enriched in the TurboID-*Cp*OGA^CD^ group relative to the TurboID-*Cp*OGA^DM^ control (log_2_ FC > 1) were regarded as O-GlcNAcylated substrates (Figure 2B). We therefore identified 336 O-GlcNAc candidate substrates from HEK293T cells (Table S1). To compare this result with known O-GlcNAc modifications, we compiled two lists of the previously identified O-GlcNAcylated proteins in HEK293T cells via either direct capture^46,47^or chemoenzymatic labeling methods^48-52^ (Table S2). Gene ontology (GO) analysis on these three datasets showed that they were enriched in similar biological processes (Figure S2E). Overlap analysis revealed that 52% (178/336) of the O-GlcNAc candidate substrates identified in our study were also present in previous reports (Figure 2D). 48 proteins were shared among the three lists (Table S3), encompassing many well-known O-GlcNAcylated proteins such as OGT, NUP153, NUP62, and HCFC1. Protein-protein interaction networks of these 48 proteins highlighted four cellular component clusters: the MLL1 complex, nuclear pores, COPII vesicle coats, and cytoplasmic stress granules (Figure 2E). Additionally, of the 158 candidate proteins that were unique in our result, 113 were annotated as O-GlcNAcylation substrates in the O-GlcNAc database (www.oglcnac.mcw.edu). These results validated that TurboID-*Cp*OGA^CD^ was able to effectively tag O-GlcNAcylated proteins with biotin for enrichment and identification.

### Region-specific O-GlcNAcylation profiling of Drosophila brain

We next generated transgenic flies harboring UAS-TurboID-*Cp*OGA^CD^ or UAS-TurboID-*Cp*OGA^DM^ via φC31 integrase-mediated site-specific recombination. To test biotinylation efficiency, we used Da-Gal4 to drive ubiquitous expression and raised the flies on biotin-containing food (100 μM) from early embryonic stage to adulthood according to previous reports^36,53^ (Figure 3A). Flies were homogenized and equal amounts of lysate were used in immunoprecipitation experiments. Similar to the result with HEK293T cells, TurboID-*Cp*OGA^CD^ catalyzed more biotinylation in the input relative to TurboID-*Cp*OGA^DM^, and more biotinylated proteins were immunoprecipitated, in which strong O-GlcNAcylation signals were detected (Figure 3B). To validate whether TurboID-*Cp*OGA^CD^ could achieve brain region-specific labeling of O-GlcNAcome with biotin tag, we selected different Gal4 to drive TurboID-*Cp*OGA^CD^ in distinct brain regions and fed the flies with biotin. Whole-mount staining of the brains showed that TurboID-*Cp*OGA^CD^ displayed specific expression patterns as expected. More importantly, staining with streptavidin-Cy3 detected strong biotinylation in the brain regions expressing TurboID-*Cp*OGA^CD^, whereas the rest of the brain showed negligible background signals (Figure 3C).

**Figure 3.**
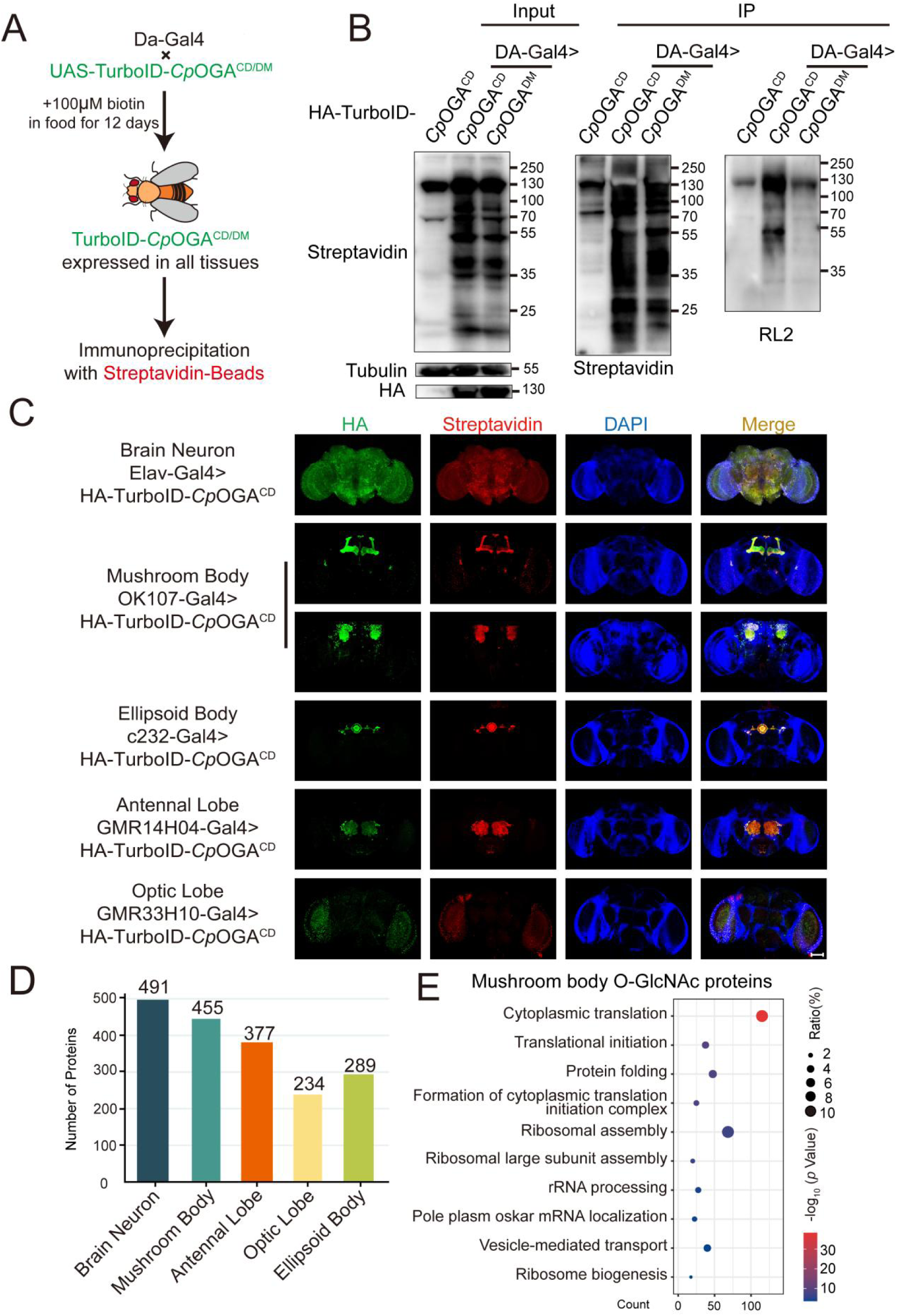
Identification of O-GlcNAc candidate substrates in different *Drosophila* brain structures using TurboID-*Cp*OGA. (A) Scheme for validating TurboID-*Cp*OGA^CD/DM^ in flies. (B) Immunoprecipitation of biotinylated proteins from flies. Biotinylation was detected by immunoblotting with streptavidin-HRP, and O-GlcNAcylation with anti-O-GlcNAc antibody (RL2). The expression of TurboID-*Cp*OGA^CD/DM^ was validated by anti-HA immunoblotting. (C) Immunostaining of *Drosophila* brains expressing TurboID-*Cp*OGA^CD^ in different brain structures. Biotinylated proteins were stained with streptavidin-Cy3 (red), and TurboID-*Cp*OGA^CD^ with anti-HA antibody. Nuclei were visualized by DAPI (blue). Scale bar: 100 μm. (D) Bar graph showing the number of O-GlcNAcylated protein candidates identified from different brain structures of *Drosophila*. (E) Gene Ontology (GO) enrichment analysis of O-GlcNAcylated protein candidates detected in the mushroom body. Bubble color indicates the -log_10_ (*p* value), and bubble size represents the ratio of genes in each category. **Figure 3—source data 1.** Raw data of all western blots from Figure 3. **Figure 3—source data 2.** Complete and uncropped membranes of all western blots from Figure 3. **Figure 3—source data 3.** Excel spreadsheet containing source data used to generate Figures 3B-E.

Subsequently, we immunoprecipitated biotinylated proteins from these fly brain lysates using streptavidin magnetic beads and performed MS analysis to identify putative O-GlcNAc substrates in different brain regions. Proteins with higher LFQ (label-free quantitation) intensity in the TurboID-*Cp*OGA^CD^ group relative to the TurboID-*Cp*OGA^DM^ control (log_2_ FC > 1 or p < 0.05) were considered as potentially O-GlcNAcylated substrates. We therefore identified 491 putative O-GlcNAcylated proteins in all neurons in fly brain (Elav-Gal4), 455 in the mushroom body (OK107-Gal4), 377 in the antennal lobe (GMR14H04-Gal4), 234 in the optic lobe (GMR33H10-Gal4), and 289 in the ellipsoid body (c232-Gal4) (Figure 3D, Table S4-S8). To obtain a functional overview of the O-GlcNAc interactome in different brain regions, GO analysis was performed to highlight the most enriched functional modules (Figure 3E, S3A-S3D). The O-GlcNAc interactome in brain neurons was enriched in chemical synaptic transmission, neurotransmitter secretion, as well as chromatin remodeling, whereas putative O-GlcNAcylated substrates in specific brain regions were involved in rather diverse biological processes, ranging from mRNA splicing to chitin-base cuticle development. Of particular interest, putative O-GlcNAcylation modifications in the mushroom body were highly clustered in processes linked to translation, including cytoplasmic translation, translational initiation, ribosome assembly, and ribosome biogenesis. To eliminate possible interference caused by varying abundance of these candidate proteins in different brain regions, we normalized the calculated O-GlcNAc level (log_2_ FC) of each substrate using its corresponding brain region-specific normalizing factor generated from the single-cell transcriptome atlas of the adult *Drosophila* brain^3^ (Figure S3E). For ease of search and use, we created an online database for tissue-<u>s</u>pecific <u>O</u>-<u>G</u>lcNAcylation <u>A</u>tlas of *Drosophila* Brain (tsOGA, http://kyuanlab.com/tsOGA/) to host these datasets (Figure S3F).

### O-GlcNAcylation affects cognitive function of Drosophila by regulating ribosomal activity in the mushroom body

The GO analysis revealed that ribosomes were enriched in the mushroom body O-GlcNAc interactome. We calculated the percentage of ribosomal components in all the proteins identified for different brain regions, and found that nearly 10% of the putative O-GlcNAc substrates in the mushroom body were from ribosomes, much higher than that in other brain regions (Figure S4A). To validate that the observed enrichment was not due to higher expression levels of these ribosomal subunits in the mushroom body, we plotted the normalized O-GlcNAc levels of the putative ribosomal substrates alongside their mRNA abundances in different brain regions. While the O-GlcNAc levels were highest in the mushroom body, their mRNA abundances were not (Figure 4A). Moreover, in the mushroom body, the O-GlcNAc levels of these ribosomal proteins showed no correlation with their mRNA abundances (Figure S4B).

**Figure 4.**
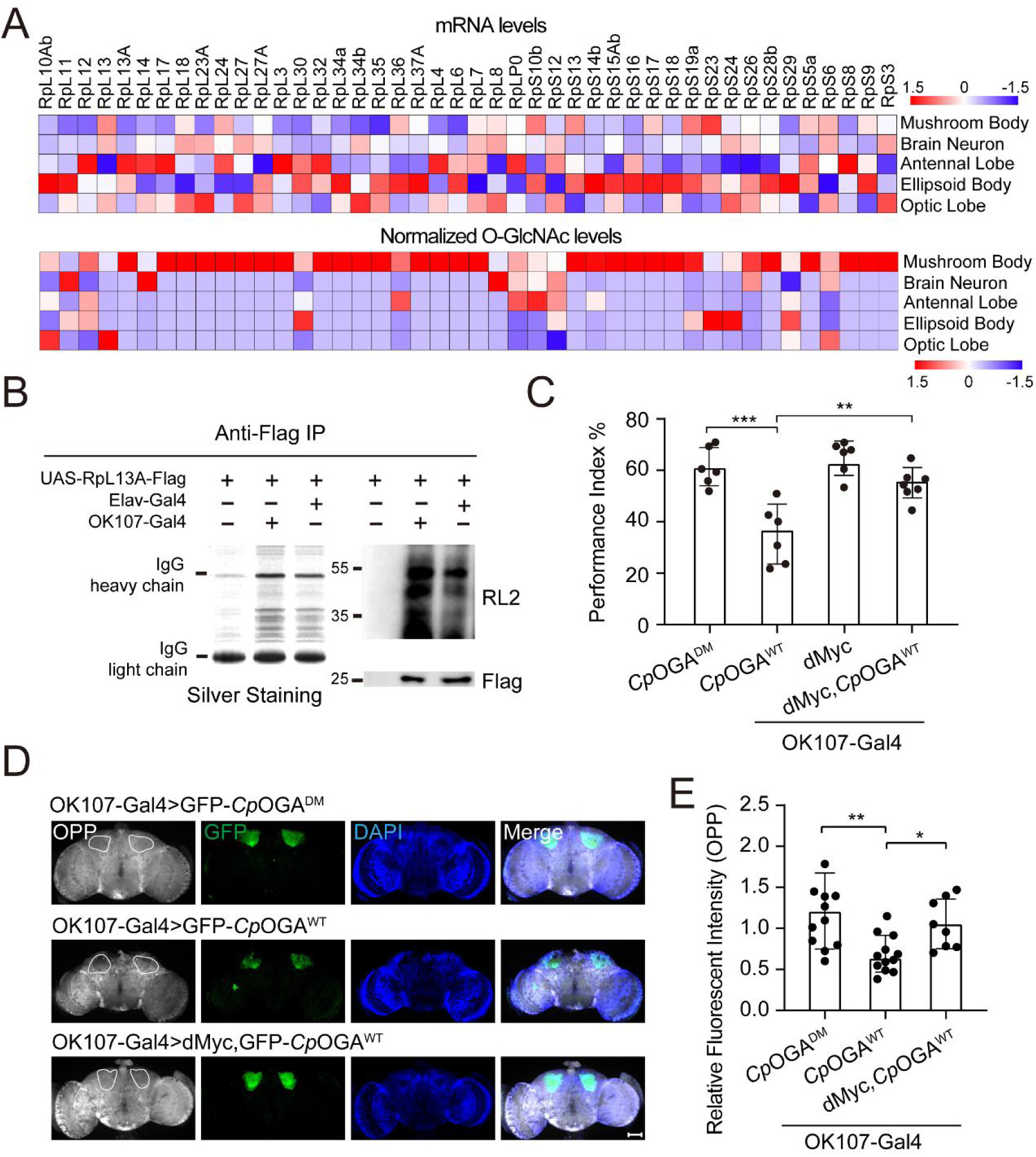
Abundant O-GlcNAcylation of ribosomal subunits in mushroom body is required for proper protein synthesis activity and olfactory learning. (A) Heatmaps showing the mRNA levels (upper) and the normalized O-GlcNAc levels (lower) of the identified ribosomal candidates in different brain regions. (B) Immunoprecipitation of ribosomes using FLAG-tagged RpL13A. The expression of RpL13A-FLAG was validated by immunoblotting with anti-FLAG antibody. Ribosomal proteins were visualized using silver staining, and O-GlcNAcylation of ribosomes was analyzed by immunoblotting with anti-O-GlcNAc antibody RL2. (C) A compilation of performance index of the indicated flies in the learning test. Learning defect of flies expressing *Cp*OGA^WT^ was corrected by selective expression of dMyc in mushroom body. Each datapoint represents an independent experiment with approximately 200 flies. (D) *Ex vivo* measurement of protein synthesis in mushroom body using the OPP assay. Brains from the indicated flies were stained with anti-GFP (green) antibody to validate *Cp*OGA expression, and OPP (grey) to quantify protein synthesis. Nuclei were visualized with DAPI (blue). Outlined areas indicate the cell bodies of kenyon cells of mushroom body. Scale bar: 100 μm. (E) Quantification of relative OPP fluorescent intensity in mushroom body regions. *p* values were determined by unpaired *t*-test, the stars indicate significant differences (\*\*\**p* < 0.001, \*\**p* < 0.01, \**p* < 0.05). Error bars represent SD. **Figure 4—source data 1.** Raw data of all western blots from Figure 4. **Figure 4—source data 2.** Complete and uncropped membranes of all western blots from Figure 4. **Figure 4—source data 3.** Excel spreadsheet containing source data used to generate Figures 4A-E.

To directly verify whether mushroom body ribosomes were hyper-O-GlcNAcylated, Flag-tagged RPL13A, a core component of the large ribosomal subunit, was expressed in brain neurons or specifically in mushroom body, driven by Elav-Gal4 or OK107-Gal4 respectively. Intact ribosomes were then isolated from these brain regions by anti-Flag immunoprecipitation^54^ (Figure 4B). Silver staining detected an array of specific bands on SDS-PAGE gel in the immunoprecipitants, indicating successful enrichment of ribosomal components. Western blot with anti-O-GlcNAc antibody RL2 showed that ribosomes purified from mushroom body contained more O-GlcNAc modifications than that from whole brain neurons. These results ascertained that ribosomal components were abundantly O-GlcNAc modified in the learning center of *Drosophila* brain.

To investigate whether high O-GlcNAcylation is required for normal ribosomal function in mushroom body, we dissected the brains from flies expressing *Cp*OGA^WT^ driven by OK107-Gal4 and measured translational activity *ex vivo* using an O-propargyl-puromycin (OPP)-based protein synthesis assay^55^ (Figure 4D). Ectopic expression of *Cp*OGA^WT^ but not the control *Cp*OGA^DM^ in mushroom body decreased local protein synthesis as visualized by the OPP fluorescent intensity (Figure 4D and 4E), suggesting that tuning down the O-GlcNAcylation compromised local ribosomal activity. Hypo-O-GlcNAcylation in mushroom body results in olfactory learning deficit (Figure 1D and 1F). We next investigated whether this cognitive phenotype was due to compromised ribosomal activity. To this end, we selected a panel of representative ribosomal components that were significantly O-GlcNAcylated in the mushroom body, and performed RNA interference (RNAi)-mediated knockdown. The RNAi induced by Da-Gal4 reduced the expression of the targeted ribosomal genes to varying degrees (Figure S4C). We then crossed the RNAi lines to OK107-Gal4 to drive specific knockdowns in mushroom body, and conducted olfactory learning assay with these flies. Downregulation of RPL11 and RPL24 in the ribosomal large subunit, and RPS3 and RPS6 in the ribosomal small subunit did not alter olfactory acuity (Figure S4E-S4F), however, they led to compromised olfactory learning ability (Figure S4D). Consequently, we reasoned that upregulation of ribosomal activity might ameliorate the cognitive defect caused by *Cp*OGA^WT^-induced hypo-O-GlcNAcylation. To test this, we increased ribosome biogenesis by overexpression of dMyc^56-58^, and observed that dMyc expression in mushroom body could restore local protein synthesis and rescue the hypo-O-GlcNAcylation induced olfactory learning deficit (Figure 4C-4E).

## Discussion

Protein O-GlcNAcylation is controlled by a very simple system consisting of only two enzymes, OGT and OGA. Yet it can dynamically modify more than 5000 protein substrates in different tissues to regulate their stability, protein-protein interactions, enzymatic activity, as well as subcellular localization upon changes of cellular metabolisms. Deciphering the spatial-temporal profiles of protein O-GlcNAcome and linking subsets of O-GlcNAc substrates to different physiological and pathological phenotypes are major obstacles in the field. In this study, we developed an O-GlcNAcylation profiling tool that allowed tissue-specific identification of O-GlcNAc candidate substrates. With this tool, we depicted the O-GlcNAc interactome in different brain regions of *Drosophila* and established an online database tsOGA (http://kyuanlab.com/tsOGA/) to facilitate future functional dissection of O-GlcNAcylation. Moreover, we consolidated a causal relationship between hypo-O-GlcNAcylation and cognitive impairment, and revealed that insufficient O-GlcNAcylation of ribosomes in the major learning center of *Drosophila* brain--the mushroom body--reduced translational activity and hence impaired the ability of associative learning.

A lot of effort has been made to identify protein O-GlcNAc modifications and many profiling methods have been established in the past 30 years. This has greatly expanded the pan O-GlcNAcome to more than 5000 substrates and boosted our knowledge on O-GlcNAcylation at the cellular level. However, given the O-GlcNAcome in different tissues and cell populations is heterogeneous and pleiotropic, our understanding of the functions of O-GlcNAc modification at organismal level remains quite limited, mainly relying on conditional knockout studies of *OGT* or *OGA*^59^. Establishment of tissue-specific landscapes of O-GlcNAc substrates in health and disease conditions is in need to fully appreciate its multifaceted functions. The strategy reported here achieved mapping the O-GlcNAcylated candidates with high spatial precision in *Drosophila* brain. With small modifications, this strategy can be readily applied to other model organisms in future studies. Moreover, using the O-GlcNAc data from different brain regions of *Drosophila*, we established a framework for a tissue-specific O-GlcNAcylation database. As more tissue-specific O-GlcNAc profiling data being generated and deposited, it will undoubtedly be a useful resource for the community to facilitate future functional interrogations of different O-GlcNAcylation substrates at organismal level.

The brain manifests high OGT expression and relies on protein O-GlcNAcylation to regulate many of its functions. Perturbed O-GlcNAcylation has been linked to neurodegenerative diseases and several key etiological factors are known O-GlcNAc substrates, such as tau^23,60^, β-amyloid (Aβ)^24^, neurofilaments (NFs)^61^, TDP-43^62^, and α-synuclein^63,64^. Particularly, O-GlcNAcylation can antagonize hyperphosphorylation of tau and stabilize it from aggregation, preventing neuronal death and tauopathies^12^. Hence, OGA inhibitors have been tested in several clinical trials to target tauopathy and early symptomatic AD, leading to a recent FDA approval of the OGA inhibitor MK-8719 as an orphan drug for tau-driven neurodegenerative disease^65^. In addition to promoting neuronal survival, O-GlcNAcylation can modify other functions of the brain. The fly model presented here shows that hypo-O-GlcNAcylation attenuates learning, and identifies proteins involved in translation as key O-GlcNAc substrates in the learning center of the fly brain. Several translational initiation factors such as eIF3, eIF4A, and eIF4G, as well as core components of ribosomes have previously been reported to harbor O-GlcNAc modifications^66-68^. But, it was not clear whether decline in global O-GlcNAcylation level influences translational activity. Our results show that formation of nascent protein polypeptides is downregulated in hypo-O-GlcNAcylated brain regions, likely resulting in the learning deficit observed. New protein synthesis is known to be required for formation and consolidation of long-term memories^69,70^. Several ribosomopathies, such as Diamond-Blackfan anemia and distal trisomy 5q, are associated with learning disabilities^71^. Our observation suggests that proper ribosomal function is indispensable for associative learning, opening up new intervention strategies for hypo-O-GlcNAcylation associated cognitive disabilities. Our O-GlcNAc profiling results also provide a rich resource for discovery of other conveyors of O-GlcNAc associated intellectual disability. For instance, the brain O-GlcNAc substrates, scu and Upf3 possess human homologues, *HSD17B10* and *UPF3B*, that are known X-linked intellectual disability risk genes^72,73^. In addition, recent studies have revealed that stress granules are tightly linked with autism spectrum disorders^74^. The enrichment of stress granule components in the O-GlcNAc substrate list suggests that O-GlcNAcylation dysregulation might be involved in autism as well. We anticipate that this study will galvanize further studies into targeting O-GlcNAcylation insufficiency to ameliorate cognitive defects commonly seen in many neurological diseases.

## Materials and Methods

### Cell cultures and generation of stable cell lines

HEK293T cells were cultured in DMEM/high glucose medium (Biological Industries, 01-052-1A) with 10% FBS (VISTECH, SE100-B) at 37 ℃ under 5% CO_2_. The *Cp*OGA^CD^ and *Cp*OGA^DM^ sequences were codon optimized to *Homo sapiens* and *Drosophila* using *Jcat*^75^. The fragments of *TurboID-CpOGA^CD^* and *TurboID-CpOGA^DM^* (*TurboID-CpOGA^CD/DM^*) were PCR amplified and cloned into pCDH-CMV-HA vectors respectively. For lentivirus preparation, HEK293T cells were transfected with *TurboID-CpOGA^CD/DM^* plasmid with the packaging plasmids pPAX2 and pMD.2G using Polyethylenimine Linear (PEI, Polysciences, 24765). The PEI-containing medium was replaced with fresh serum-containing DMEM medium after 8 h, and the viral supernatants were collected 48 h and 72 h post-transfection. The viral supernatants were centrifuged at 10000 g for 1 h at 4 ℃, and the pellet was dissolved in PBS (Biological Industries, 02-023-1A). HEK293T cells were infected in 6-well plates and selected with 1 µg/mL Puromycin (Selleck, s7417) in the medium for at least 5 d. For biotin labeling, the TurboID-*Cp*OGA^CD^ or TurboID-*Cp*OGA^DM^ expressing HEK293T cells were labeled with 10 to 100 µM biotin (Merck, B4501) in the medium for 15 min to 3 h. Labeling was stopped by placing cells on ice and washing cells three times with PBS (Biological Industries, 02-023-1A).

### Drosophila Stocks and Genetics

All flies were raised on standard fly food at 25 °C. Biotin food was prepared by adding 200 mM biotin (Merck, B4501) to hot (∼60 ℃) standard fly food and dissolved to a final concentration of 100 μM^53^. The strains used in this study were as follow: *w1118*, *;sco/cyo;TM3/TM6B*, *Da-Gal4* (Gift from Kun Xia’s lab), *Elav-Gal4* (Gift from Zhuohua Zhang’s lab), *OK107-Gal4*, *201Y-Gal4* (Gift from Ranhui Duan’s lab), *C232-Gal4* (BDSC, #30828), *GMR14H04-Gal4* (BDSC, #48655), *GMR33H10-Gal4* (BDSC, #49762), *Tub-Gal80*^ts^, *uas-RPL13A-FLAG*, *uas-dMyc* (Gift from Jun Ma’s lab), *uas-shLuciferase* (Gift from Zhuohua Zhang’s lab), *uas-shRPL5* (THU0670), *uas-shRPs26* (THU0747), *uas-shRPL24* (THU1411), *uas-shRPS6* (THU0864), *uas-shRPL11* (TH201500769.S), *uas-shRPS3* (THU1958), *uas-shRPL32* (TH201500773.S), *uas-shRPS28b* (THU1037). Our study established two transgenic fly lines (*UAS-HA-TurboID-CpOGA^CD^* and *UAS-HA-TurboID-CpOGA^DM^*). *TurboID-CpOGA^CD/DM^* fragments were cloned into pUASz-HS-HA vectors respectively using Gibson assembly (NEB). Constructs with the attB sequence were injected into flies (*y1, w67c23; P(CaryP) attP2*) to initiate the φC31 integrase-mediated site-specific integration (UniHuaii). The resulted adult flies (G0) were crossed to double balancer to get the F1 generations.

### Olfactory learning and memory

Behavioral experiments were carried out in an environmental chamber at 25 °C and 70% humidity as previously described^37^. We tested the acuity of flies against two aversive odors, 4-methylcyclohexanol (MCH, Sigma, 104191) and 3-octanol (OCT, Sigma, 218405). Approximately 100 flies were placed in the center compartment of the T-maze, where the collection tubes were snapped into place at the choice point and the air and aversive odor tubes were connected with the distal ends of the collection tubes. Flies were allowed to choose between air versus aversive odor for 2 min. After the choice period, the sliding center compartment was pulled up quickly, trapping the flies in the collection tubes they had chosen. Flies in each collection tube were anesthetized and counted. Performance index (PI^odor^) was determined as the number of flies in the air side (n(Air)) minus the number in the aversive odor side (n(odor)) divided by the total number of flies (n(Air)+n(odor)) and multiplied by 100%.

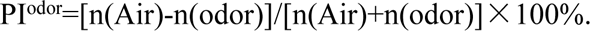

If the experimental group flies have similar odor avoidance to that of control, they will be used for subsequent olfactory learning test.

After confirming that the flies to be tested have avoidance behavior in response to electric shock, flies were trained to associate an aversive odor (MCH or OCT) used as a conditioned stimulus (CS) with electric shock. The experiment comprised two phases: the flies were trained in the first phase, and the trained flies were tested in the second phase. During training, approximately 100 flies were simultaneously exposed to odor 1 (CS^+^) and electric shock (60 V) in a training tube for 1 min. Then, they were exposed to the blank odor (air) for 1 min before receiving odor 2 (CS^-^) without electric shock for 1 min, followed by the blank odor (air) for 1 min. Immediately after training, flies were transferred to the central chamber of the T-maze and retained there for 1 min. To measure learning, The center chamber was slid smoothly into register with the choice point of the T-maze and the MCH and OCT odor tubes were supplied from the two distal ends of the collection tube to let the flies choose between the two odors for 2 min. The central chamber then was pulled up quickly, trapping the flies in the collection tube they had chosen. Flies in each collection tube were anesthetized and counted. We calculated the Performance Index (PI) for each condition as the number of flies avoiding the shock-paired odor (CS^-^) minus the number of flies choosing the shock-paired odor (CS^+^) divided by the total number of flies (CS^-^ + CS^+^) and multiplied by 100%.

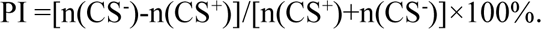

In each experiment, we calculated the mean PI from two trials: one in which MCH was the shock-paired odor, and the other in which OCT was the shock-paired odor. This method removed any potential bias caused by the flies having a stronger preference for one odor over the other. Therefore, each point in the bar graph consisted of approximately 200 flies (male: female = 1:1), with half of the flies trained to one odor, and the other half trained to the other odor.

For the temporally controlled *Cp*OGA expression in adult mushroom body, the flies were initially maintained at 19 ℃ until adulthood. Then, the flies were transferred to 29 ℃ for 3-5 d to inactivate Gal80^ts^ and hence allow the expression of *Cp*OGA. The behavioral experiments were carried out subsequently.

### Western blot assay

The HEK293T cells and flies were lysed in lysis buffer (2% SDS, 10% glycerol, and 62.5 mM Tris-HCl, pH 6.8) supplemented with protease inhibitor cocktail (1:100, Sigma, P8340), and PMSF (1:100, Sigma, P7626) and 50 µM Thiamet-G (Selleck, s7213). Lysates were clarified by centrifugation at 13000 rpm for 30 min at 4 ℃, and the protein concentration was determined using BCA assay (Beyotime, p0009). Proteins were mixed with an equal volume of SDS sample buffer (2% β-Mercaptoethanol) and boiled for 10 min at 95 ℃. Proteins were separated by 10% SDS-PAGE (90 V, 30 min; 120 V, 1 h) and transferred to a Polyvinylidene Fluoride (PVDF, Millipore, IPVH00010) membrane (290 mA, 90 min). The PVDF membrane was blocked with 5% non-fat milk for 1 h, then incubated with primary antibodies overnight at 4 ℃, and then incubated with secondary antibodies (1:5000, Thermo Fisher Scientific) for 1 h at room temperature. The signal was detected using ECL substrates (Millipore). Primary antibodies were dissolved in 5% BSA (Biofroxx, 4240GR005) and the dilutions were: Streptavidin-HRP (1:2000, GenScript, M00091), RL2 (1:1000, Abcam, ab2739), HA (1:3000, Cell Signaling Technology, 3724), Tubulin (1:3000, Cell Signaling Technology, 12351S), FLAG (1:3000, Cell Signaling Technology, 14793). For the Western blot experiment in Figure S2C and D, cells were cultured in the medium supplemented with 25 μM Thiamet-G (Selleck, s7213) or 25 μM OSMI-1(Sigma, SML1621) for 6 h before lysis. For the experiment in Figure 4D, the gel was stained with Fast Silver Stain Kit (Beyotime, P0017S).

### Immunoprecipitation

For the immunoprecipitation experiment in Figure 2C and 3B, the HEK293T cells (1×10^7^ cells per sample) and flies (∼20 flies per sample) were lysed in RIPA lysis buffer (50 mM Tris pH 8.0, 150 mM NaCl, 0.1% SDS, 0.5% Sodium deoxycholate, 1% NP40, 10 mM NaF, 10mM Na_2_VO_4_, 50 µM Thiamet-G) supplemented with protease inhibitor cocktail (1:100, Sigma, P8340) and PMSF (1:100, Sigma, P7626) on ice for 30 min. After centrifugation at 13000 g for 30 min at 4 ℃, the supernatants were transferred to new tubes. The protein concentration was determined using BCA assay (Beyotime, p0009). Streptavidin magnetic beads (MCE, HY-K0208) were washed twice with RIPA lysis buffer, and incubated with the same amount of lysate from TurboID-*Cp*OGA^CD^ or control samples on a rotator overnight at 4 ℃. The beads were washed twice with 1 mL of RIPA lysis buffer, once with 1 mL of 1 M KCl, once with 1 mL of 0.1 M Na_2_CO_3_, once with 1 mL of 2 M urea in 10 mM Tris-HCl (pH 8.0), and twice with 1 mL RIPA lysis buffer. After that, the beads were resuspended in SDS sample buffer and boiled for 10 min at 95 ℃. Finally, samples were stored at −80 ℃ for future analysis.

The immunoprecipitation experiment in Figure 4B was performed as previously described^54^. Briefly, fly brains (∼40 fly brains per sample) were lysed in ribo-lysis buffer (50 mM Tris-HCl pH 7.4, 12 mM MgCl_2_, 100 mM KCl, 1 mM DTT, 1% NP-40, 100 µg/mL cycloheximide, 50 µM Thiamet-G) supplemented with protease inhibitor cocktail (1:100, Sigma, P8340) and PMSF (1:100, Sigma, P7626) on ice for 30 min. After centrifugation at 13000 g for 30 min at 4 ℃, the supernatants were transferred to new tubes. The protein concentration was determined using BCA assay (Beyotime, p0009). Anti-FLAG M2 affinity gels (Sigma, A2220) were washed twice with ribo-lysis buffer, and incubated with tissue lysates on a rotator overnight at 4 ℃. The beads were washed three times with 1 mL of high salt buffer (50 mM Tris-HCl pH 7.4, 12 mM MgCl_2_, 300 mM KCl, 1 mM DTT, 1% NP-40, 100 µg/mL cycloheximide). The beads were resuspended in SDS sample buffer and boiled for 10 min at 95 ℃. Finally, samples were stored at −80 ℃ for future analysis.

### Immunofluorescence

The adult fly brains were dissected in PBS and fixed with 4% paraformaldehyde (PFA, Biosharp, BL539A) for 1 h at room temperature. The brains were washed three times with PBS (Biological Industries, 02-023-1A) and then permeabilized and blocked in 5% BSA (Biofroxx, 4240GR005) in 0.3% PBST (PBS with 0.3% Triton X-100) for 90 min at room temperature. After being washed three times with PBS, the brains were incubated with primary antibodies overnight at 4 ℃, washed three times with PBS, and incubated with secondary antibodies (1:200, Thermo Fisher Scientific) and DAPI (1:500, Sigma, D9542) for 1 h at room temperature. The brains were then washed three times with PBS and imaged by confocal fluorescence microscopy (Zeiss LSM880) with a 20× objective. Z-stacks were acquired with a spacing of 1 μm. Primary antibodies were dissolved in 5% BSA (Biofroxx, 4240GR005) and the dilutions were: Streptavidin-Cy3 (1:200, BioLegend, 405215), RL2 (1:200, Abcam, ab2739), HA (1:200, Cell Signaling Technology, 3724) and GFP (1:200, Cell Signaling Technology, 2955).

### Measurement of Protein Synthesis

The protein synthesis in fly brains was assessed using the Click-iT Plus OPP Alexa Fluor® 594 Protein Synthesis Assay Kit (Thermo Fisher Scientific, C10457). Fly brains were dissected in *Drosophila* medium (Gibco, 21720024) and then incubated in medium containing 1:1000 (20 µM) of Click-iT OPP reagent at room temperature for 30 min. The brains were washed three times with PBS, and then fixed with 4% PFA (Biosharp, BL539A) for 1 h at room temperature. The brains were permeabilized and blocked in 5% BSA (Biofroxx, 4240GR005) in 0.3% PBST (PBS with 0.3% Triton X-100) for 90 min at room temperature, and then washed three times with PBS. The brains were incubated with primary antibodies (GFP, 1:200, Cell Signaling Technology, 2955) overnight at 4 ℃, washed three times with PBS, and incubated with secondary antibodies (1:200, Thermo Fisher Scientific) and DAPI (1:500, Sigma, D9542) for 1 h at room temperature. For the Click-iT reaction, brains were incubated in the Click-iT reaction cocktail in the dark at room temperature for 30 min. Brains were then washed three times with PBS and imaged by confocal fluorescence microscopy (ZEISS LSM880).

### RT-qPCR

RNA was extracted from flies using TRIzol (Life Technologies, 87804), and 1 μg total RNA was reverse transcribed to generate cDNA using RevertAid First Strand cDNA Synthesis Kit (Thermo Fisher Scientific, K1621). The cDNA was then used as templates and qPCR was performed using the SYBR Green qPCR Master Mix (Solomon Biotech, QST-100) on the QuantStudio3 Real-Time PCR system (Applied Biosystems). The expression levels for each gene were normalized to Actin. Detailed information about the primers was listed in Table S9.

### Protein Identification by LC-MS/MS

The HEK293T cells (2 × 10^7^ cells per sample) and fly brains (∼200 fly brains that expressed TurboID-*Cp*OGA^CD/DM^ in brain neurons per sample, ∼800 fly brains that expressed TurboID-*Cp*OGA^CD/DM^ in other brain structures per sample, three biological replicates) were immunoprecipitated with streptavidin magnetic beads as described above. The supernatants were used for SDS-PAGE separation and minimally stained with Coomassie brilliant blue (Solarbio, C8430-10g). The gels were cut into small pieces, and reduced and alkylated in 10 mM DTT and 55 mM IAA (Merck, I6125) respectively. For digestion, 0.5 µg sequencing-grade modified trypsin was added and incubated at 37 ℃ overnight. The peptides were then collected, desalted by StageTip (Thermo Fisher Scientific, 87782) and resolved in 0.1% formic acid before analysis by mass spectrometry. Mass spectrometry analysis was performed using Q Exactive HF-X mass spectrometer (Thermo Fisher Scientific) coupled with Easy-nLC 1200 system. Mobile phases A and B were water and 80% acetonitrile, respectively, with 0.1% formic acid. Protein digests were loaded directly onto an analytical column (75 µm × 15 cm, 1.9 µm C18, 1 µm tip) at a flow rate of 450 nL/min. Data were collected in a data-dependent manner using a top 25 method with a full MS mass range from 400 to 1400 m/z, 60,000 resolutions, and an AGC target of 3 × 10^6^. MS2 scans were triggered when an ion intensity threshold of 4 × 10^5^ was reached. A dynamic exclusion time of 30 sec was used. Ions with charge state 6-8 and more than 8 were excluded.

### Data analysis

The raw data were imported into the MaxQuant software to identify and quantify the proteins. The following parameters were used: trypsin for enzyme digestion; oxidation of methionine, acetylation of the protein N terminus, biotinylation of lysine and protein N terminus and HexNAc (ST) as variable modifications; carbamidomethyl (C) as fixed modification. We used the canonical human protein database (containing 20379 reviewed protein isoforms) or *Drosophila melanogaster* protein database (containing 22088 protein isoforms, including reviewed and unreviewed sequences) for database searching separately. The false discovery rate (FDR) was 1% for peptide-spectrum matches (PSM) and protein levels. For the proteomics data of different brain regions of *Drosophila*, we used label-free quantitation (LFQ) to determine the relative amounts of proteins among 3 replicates. Perseus software was used to filtered out all contaminates identified by MaxQuant (contaminant proteins, reversed proteins, proteins only identified by site). A pseudo-count of 1 was added to protein intensities in order to avoid taking log of 0. We generated log_2_ Fold Change (log_2_ FC) values for each protein in the TurboID-*Cp*OGA^CD^ group relative to the TurboID-*Cp*OGA^DM^ control. For the proteomics data of HEK293T cell, only proteins identified with at least 2 peptides were considered for further analysis. Proteins were considered as O-GlcNAcylated substrates when differences in log_2_ FC of TurboID-*Cp*OGA^CD^ group with relative to the TurboID-*Cp*OGA^DM^ control were higher than 1. For the proteomics data from different brain regions of *Drosophila*, only proteins identified with at least 2 peptides and in at least 2 of the 3 replicates of TurboID-*Cp*OGA^CD^ were included for further analysis. A two-tailed unpaired student’s t-test was applied in order to determine the statistical significance of the differences. Proteins were considered as O-GlcNAcylated substrates when differences in log_2_ FC of TurboID-*Cp*OGA^CD^ group with relative to the TurboID-*Cp*OGA^DM^ control were higher than 1 or statistically significant (*p* < 0.05).

To adjust the intereference caused by varying abundance of the putative O-GlcNAc substrates in different brain regions, single-cell transcriptomic data of the entire adult *Drosophila* brain (GEO: GSE107451) ^3^ was used to generate a normalizing factor for each substrate. Briefly, the annotated cell clusters were categorized into different brain regions. Then, the average mRNA expression level of each gene within a certain brain region was calculated. The normalizing factor was defined as the ratio of the average mRNA expression level of a given gene in neurons from a specific brain region to the average mRNA expression level of the same gene in neurons from the whole brain (Table S10). The normalized O-GlcNAc level was generated as the O-GlcNAc level (log_2_ FC) of a putative O-GlcNAcylated protein divided by its normalizing factor in a certain brain region (Table S11).

### Website

The website was created to browse through the O-GlcNAc database (www.kyuanlab.com/tsOGA), using the database management system Centos and the Uwsgi web framework. Backend servers were developed by Python programming language (version 3.7). GNU/Linux Debian-based systems with gunicorn (Python http) and NginX were used for development and production of the website. The website search function was based on MySQL database.

### Quantification and Statistical Analysis

To quantify fluorescent intensities in different *Drosophila* brain regions, whole brain images were stitched together using the stitching algorithm in ZEN software (Zeiss), and maximum intensity projection was produced. The images were then analyzed using ImageJ software. Mean fluoresecent intensity of the whole brain or ROI was measured, and the relative fluorescent intensity was calculated as a ratio of the mean fluorescent intensity in ROI to that of the whole brain.

GO enrichment analyses of O-GlcNAcome in HEK293T cells and *Drosophila* were performed using *DAVID*. Protein-protein interaction (PPI) network of O-GlcNAcome in HEK293T cells was performed using *STRING*. GraphPad Prism was used for statistical analysis and the student’s t-test was used to determine statistical significance. Bubble plots, pie plots and bar graphs were created using *Hiplot*, venn plots were created using *jvenn*.

## Data and materials availability

### Data availability

The accession numbers for the mass spectrometry data were PXD040547 and PXD040412 on the ProteomeXchange Consortium PRIDE partner repository.

### Materials availability

All cells and fly strains generated in this study are available upon request to the lead contact (see above).

### Lead contact

Further information and requests for resources and reagents should be directed to and will be fulfilled by the lead contact, Dr. Kai Yuan.

## Supplemental Table

Table S1. O-GlcNAcylated proteins identified by TurboID-*Cp*OGA^CD^ from HEK293T cells.

Table S2. Previously identified O-GlcNAcylated proteins from HEK293T cells, related to Figure 2D and S2E.

Table S3. 48 proteins shared among the three datasets, related to Figure 2D and Figure 2E.

Table S4. O-GlcNAcylated proteins identified by TurboID-*Cp*OGA^CD^ from brain neuron of *Drosophila*.

Table S5. O-GlcNAcylated proteins identified by TurboID-*Cp*OGA^CD^ from mushroom body of *Drosophila*.

Table S6. O-GlcNAcylated proteins identified by TurboID-*Cp*OGA^CD^ from antennal lobe of *Drosophila*.

Table S7. O-GlcNAcylated proteins identified by TurboID-*Cp*OGA^CD^ from ellipsoid body of *Drosophila*.

Table S8. O-GlcNAcylated proteins identified by TurboID-*Cp*OGA^CD^ from optic lobe of *Drosophila*.

Table S9. Sequences of all the primers used in this study.

Table S10. Cell clusters in different brain regions generated from single-cell transcriptomic data.

Table S11. The normalized O-GlcNAc levels of O-GlcNAcylated proteins in different brain regions.

## Acknowledgments

We gratefully acknowledge Drs. Jilong Liu, Hai Huang, Feng He, Yan Chen, Pishun Li, Ranhui Duan, the Developmental Studies Hybridoma Bank, the Bloomington *Drosophila* Stock Center, and TsingHua Fly Center for reagents and fly stocks. We thank colleagues in the center for medical genetics and members of the Yuan lab for helpful discussions. This project has been supported by the National Natural Science Foundation of China (grants 92153301, 91853108, and 32170821 to K.Y, 32101034 to F.C), Department of Science & Technology of Hunan Province (grants 2021JJ10054 and 2019SK1012 to K.Y), Central South University (2021zzts0566 to H.Y, 2019zzts046 to Y.Z, and the innovation-driven team project 2020CX016), a Novo Nordisk Fonden Laureate award (NNF21OC0065969) and a Villum Fonden Investigator (00054496) to D.M.F.v.A.

## Author contributions

Conceptualization: K.Y. and D.M.F.v.A.; Methodology: H.Y., D.L., Y.Z., S.M., F.C., H.Q., B.Y., K.Y.; Validation: Y.Z., R.T., X.F., F.C., L.L.; Software: H.Y., S.M.; Formal Analysis: H.Y., Y.Z., K.Y.; Investigation: H.Y., D.L., Y.Z., S.M., K.Y.; Resources: H.Q., Z.Z., D.M.F.v.A., B.Y., K.Y.; Data Curation: H.Y., D.L., S.M.; Writing-Original Draft: H.Y.; Writing-Review & Editing: D.M.F.v.A., K.Y.; Visualization: H.Y., Y.Z., S.M., K.Y.; Supervision: Z.Z., K.Y.; Project Administration: K.Y., L.L.; Funding Acquisition: K.Y.

---

**Figure S1.**
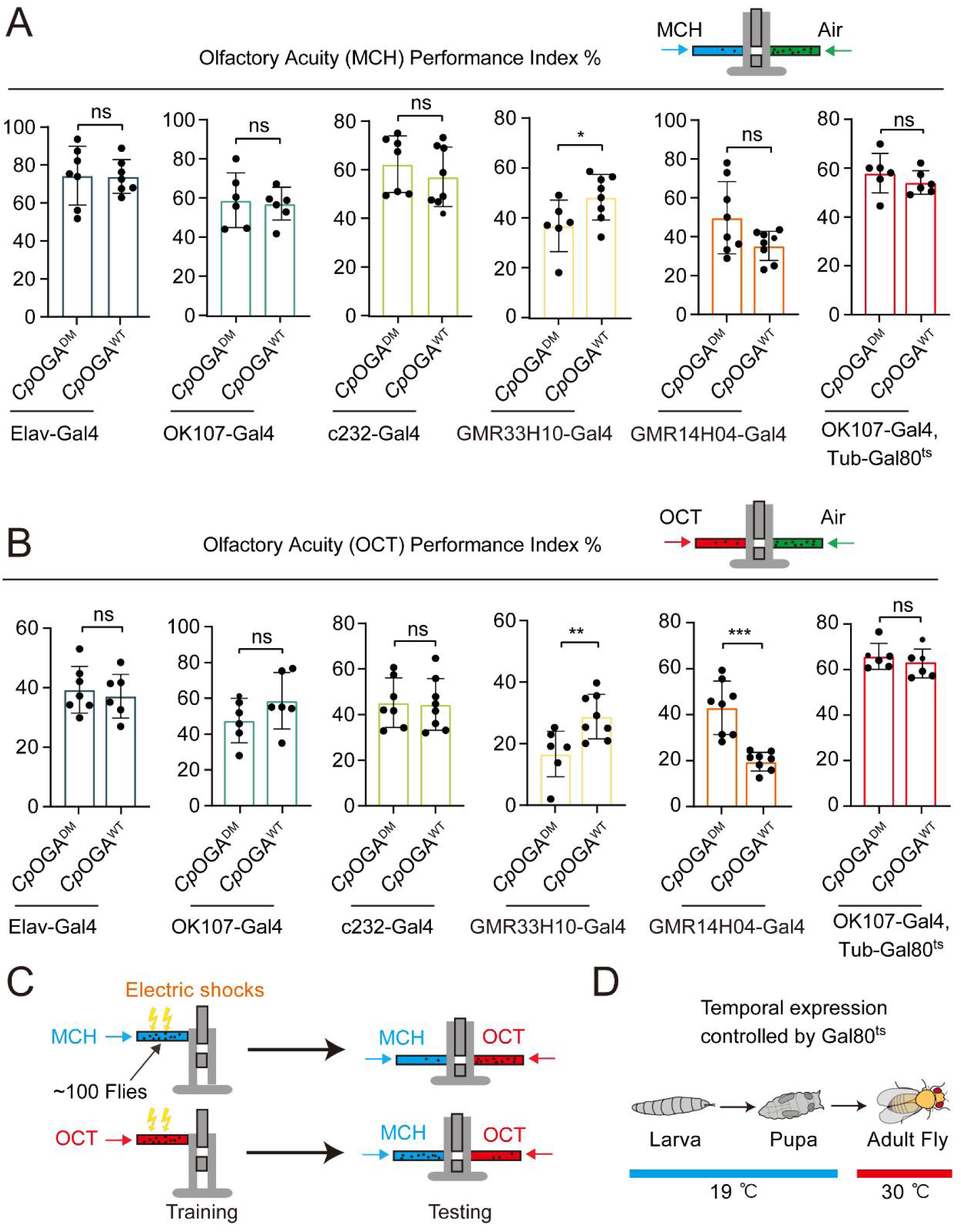
Impacts of reduction of O-GlcNAcylation in different brain structures on odor acuity towards MCH or OCT. (A-B) Bar graphs showing the odor acuity performance index of flies expressing *Cp*OGA^WT^ or *Cp*OGA^DM^ in the indicated brain regions. *p* values were determined by unpaired *t*-test, the stars indicate significant differences (\*\*\**p* < 0.001, \*\**p* < 0.01, \**p* < 0.05, and ns, not significant, *p* ≥ 0.05). Error bars represent SD. (C) Schematic of *Drosophila* learning test. Black spots represent flies. In one trial, about 100 flies were trained to associate one of the two aversive odors (MCH or OCT) with electric shock on the upper part of a T-maze, and tested for odor preference on lower part of the T-maze. Each independent experiment consisted of two trials with different odors coupled to the electric shock. The mean PI was calculated and plotted as one datapoint. (D) Temperature shifting scheme for Gal80^ts^ restricted expression of *Cp*OGA till adulthood. **Figure S1—source data 1.** Excel spreadsheet containing source data used to generate Figures S1A-B.

**Figure S2.**
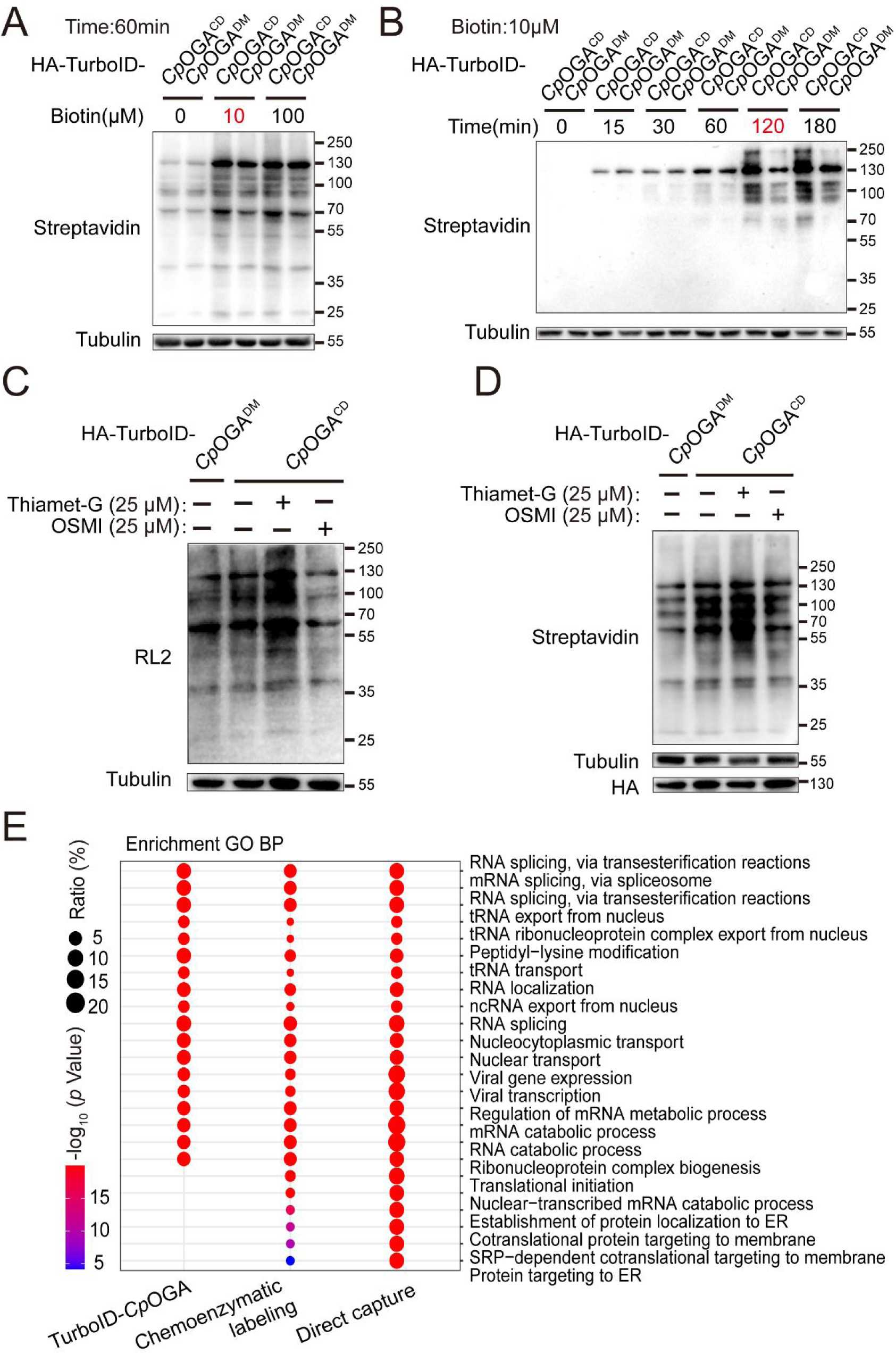
Validation and optimization of TurboID-*Cp*OGA^CD^ mediated intracellular labeling. (A-B) Determination of the optimal biotin concentration and incubation time for TurboID-*Cp*OGA^CD^ labeling in HEK293T cells. Biotinylated proteins were detected by immunoblotting with streptavidin-HRP. (C-D) Immunoblotting of cells treated with Thiamet-G or OSMI-1 using anti-O-GlcNAc antibody (RL2) or streptavidin-HRP. The expression of TurboID-*Cp*OGA^CD/DM^ was detected by anti-HA immunoblotting. (E) Bubble plot showing the Gene Ontology (GO) enrichment analysis of candidate O-GlcNAcylated substrates identified by the indicated methods. Bubble color indicates the -log_10_ (*p* value), and bubble size represents the ratio of genes in each category. **Figure S2—source data 1.** Raw data of all western blots from Figure S2. **Figure S2—source data 2.** Complete and uncropped membranes of all western blots from Figure S2. **Figure S2—source data 3.** Excel spreadsheet containing source data used to generate Figures S2A-E.

**Figure S3.**
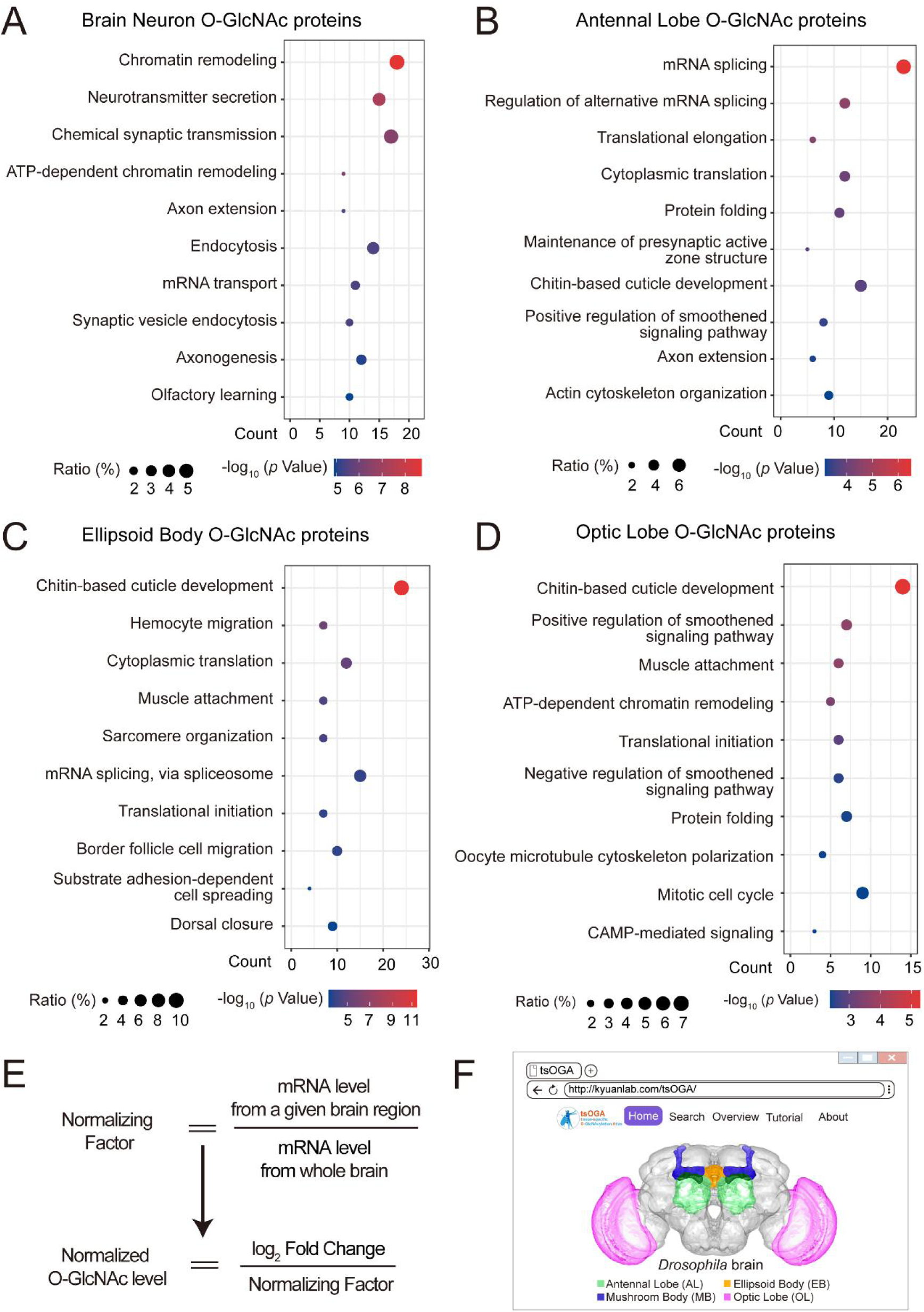
GO analysis of candidate O-GlcNAc substrates from different brain regions of *Drosophila*. (A-D) Gene Ontology (GO) enrichment analysis of potentially O-GlcNAcylated proteins in whole brain neurons (A), antennal lobe (B), ellipsoid body (C), and optic lobe (D). Bubble color indicates the -log_10_ (*p* value), and bubble size represents the ratio of genes in each category. (E) Strategy to normalize the O-GlcNAcylated protein level with expression level for each candidate substrate in different brain regions. A normalizing factor for each substrate in a given brain region was calculated using single cell RNA-seq expression data, and the adjusted O-GlcNAc level was determinted as the log_2_ FC divided by its normalizing factor. (F) The front-page of the tsOGA (tissue-specific O-GlcNAcylation Atlas of *Drosophila* brain) website. **Figure S3—source data 1.** Excel spreadsheet containing source data used to generate Figures S3A-D.

**Figure S4.**
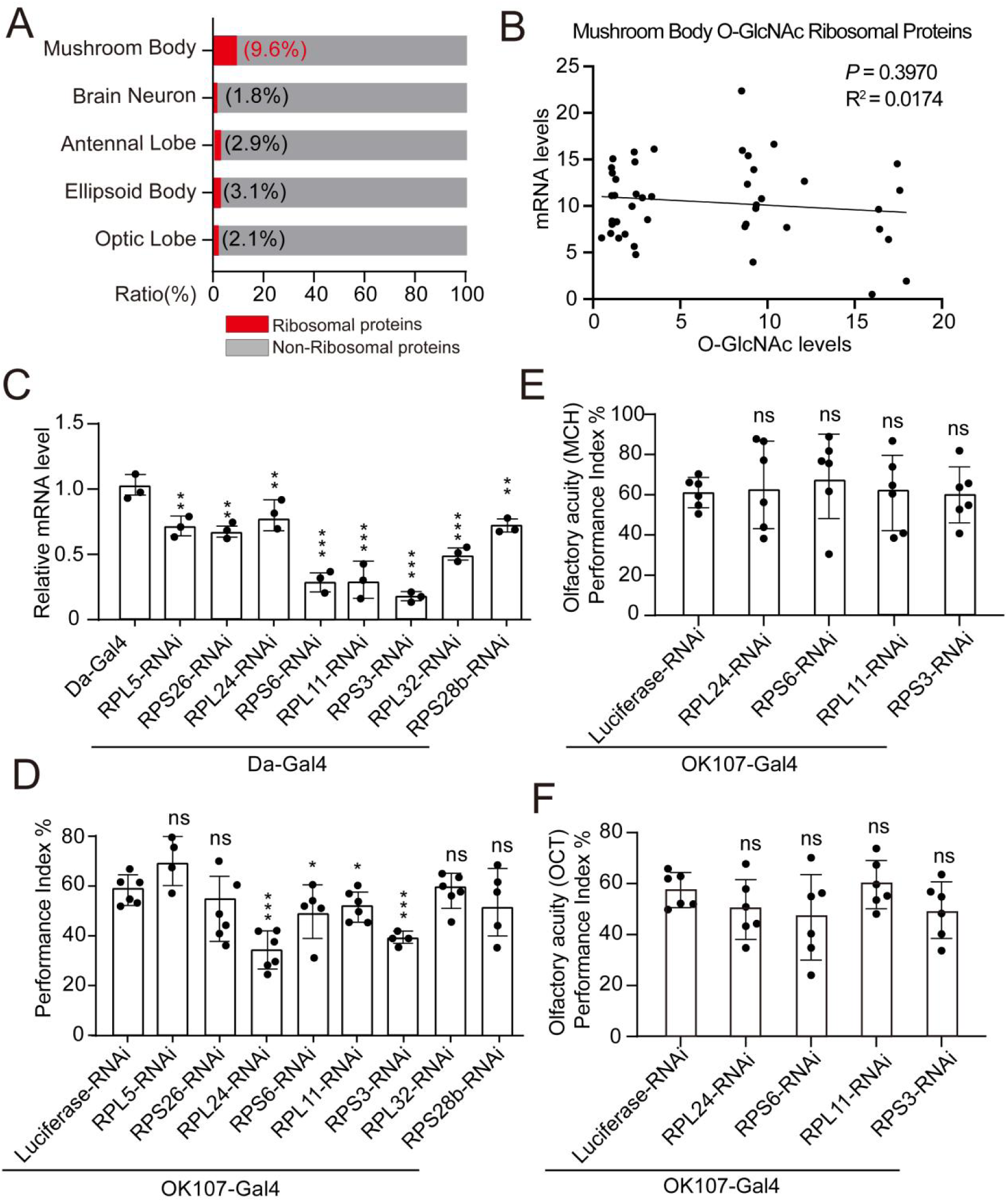
Weakened ribosomal activity in mushroom body impacts olfactory learning. (A) Bar chart representing the proportion of ribosomal proteins in the O-GlcNAcome of different brain structures identified by TurboID-*Cp*OGA. (B) Correlation analysis between mRNA levels and O-GlcNAc levels of candidate ribosomal substrates in mushroom body. Linear regression analysis was performed and no significance was noted (*p* ≥ 0.05). (C) qPCR analysis of the indicated ribosomal components expression after shRNA-mediated knockdown. (D) A compilation of performance index of the control and ribosomal subunits knockdown flies in the learning test. Each datapoint represents an independent experiment with approximately 200 flies. (E-F) Bar graphs showing the odor acuity performance index of the control and ribosomal subunits knockdown flies. *p* values were determined by unpaired *t*-test, the stars indicate significant differences (\*\*\**p* < 0.001, \*\**p* < 0.01, \**p* < 0.05, and ns, not significant, *p* ≥ 0.05). Error bars represent SD. **Figure S4—source data 1.** Excel spreadsheet containing source data used to generate Figures S4A-F.

